# The individual brain: mapping variability in hemispheric functional organization across cognitive domains in a representative sample

**DOI:** 10.1101/2025.11.05.686697

**Authors:** Emma M. Karlsson, Robin Gerrits, Guy Vingerhoets

## Abstract

Several core cognitive networks in the human brain show marked left–right differences in their functional organization. While these asymmetries are well described at the level of individual functions, overarching patterns of variability in hemispheric functional organization across multiple domains have not been mapped in a representative sample. To address this gap, we conducted a large-scale neuroimaging study with 200 participants (100 left-handers and 100 right-handers) to map hemispheric phenotypes across four distinct cognitive domains: language, tool use, spatial attention, and face perception. We challenge the traditional one-size-fits-all view of hemispheric organization by showing that deviations from the typical pattern of functional segregation are more prevalent than generally assumed, in both left- and right-handers. As predicted, variation in asymmetry was more pronounced in the left-handed sample. Critically, we found no evidence that the prototypical, “textbook”, pattern of brain organization confers any advantage in general cognitive ability or IQ. These results challenge the assumption of a single optimal brain organization and demonstrate that hemispheric functional segregation in humans is much more variable than anticipated.

## Background

Hemispheric functional specialization is one of the major organizational principles in the brain. Language^1,2^ and tool-related knowledge^3–5^, for example, rely predominantly on left-hemispheric networks in the majority of the population. By contrast, the right hemisphere plays a dominant role in spatial attention^6,7^ and face processing^8,9^. Mapping these prototypical patterns of functional segregation has been central to understanding human brain organization. Yet because each function is typically examined in isolation, we know far less about the relationships between different asymmetrical functions within individuals.

The assumption that most people share the aforementioned *prototypical* hemispheric organization is deeply ingrained in neuroscience and is predominantly established on data averaged across individuals. Although case reports of atypical hemispheric patterns have appeared for more than a century, only in recent years have these lateralization phenotypes been recognized as essential for understanding the full breadth of human brain variability^10–13^. It is well-established that some individuals have one or more functions shifted to the opposite hemisphere, e.g., language in the right hemisphere or face processing in the left, whereas in others they are distributed more symmetrically across both hemispheres. Historically, such deviations from typical dominance have been considered rare and were most often associated with left-handedness^1^, or with neurological and neurodevelopmental conditions such as schizophrenia or developmental dyslexia^14^, rather than with natural variability in asymmetry. This assumption has, in turn, led to the routine exclusion of left-handers from thousands of neuroimaging studies^15,16^, likely underestimating the true diversity of human brain organization.

The tendency to associate atypical hemispheric organization with left-handedness also means that people often overlook the fact that hemispheric variability is also present among right-handers. Emerging evidence challenges the idea of a uniform, “*textbook*” pattern of hemispheric specializations in this group. For example, although only about 5% of right-handers show atypical, right-hemispheric dominance for language^1,2^, face perception more frequently lateralizes to the atypical hemisphere^17–19^. These observations also challenge the view of strict hemispheric complementarity^20–22^, which assumes that a reversal in hemispheric dominance for one function entails a reversal for all functions, and underscore that interindividual variability in hemispheric asymmetry is not just an exception confined to the left-handed minority group. Capturing the full extent of this variability requires systematic mapping of hemispheric organization across individuals. As neuroscience increasingly emphasizes the importance of variability in brain structure and function, closing this gap in the literature is vital for advancing theoretical models of brain organization.

Beyond describing variability, a central challenge is understanding why the brain typically develops asymmetrically and why most asymmetries have strong population biases to one hemisphere. Despite the fundamental role of hemispheric specialization, its origins are poorly understood, and surprisingly few theories address why asymmetry evolved. Those that do often assume that deviations from *prototypical segregation* carry a cost, with typical asymmetry patterns viewed as an evolutionary solution for optimal neural processing^23–26^. If this configuration indeed reflects functional optimization, atypical asymmetry may be biologically disadvantageous, forcing otherwise segregated functions to share neural or computational resources. Thus, one of the core questions in the broader research framework on the advantages and disadvantages of hemispheric asymmetries is whether specific patterns of asymmetry influence cognitive task performance.

Although several studies have examined the relationship between functional lateralization and cognitive performance, results have been inconsistent and occasionally contradictory^27–29^. One key limitation that may underlie these mixed findings is that most studies have focused on a single lateralized function in isolation. However, emerging evidence indicates that understanding hemispheric organization requires considering how multiple asymmetries co-segregate within the same individual^11,12,30,31^. For example, Quin-Conroy et al.^12^ reviewed 17 studies on language and spatial attention lateralization, two functions that typically reside in opposite hemispheres, and found preliminary evidence that co-locating these functions in the same hemisphere may impair cognitive abilities, such as language comprehension, spatial skills, and reading. In a previous study in the lab, we also found that left-handed individuals with high deviation from typical functional segregation performed worse on general cognitive tests^30^. Together, these findings suggest that prototypical hemispheric segregation may confer cognitive advantages, whereas atypical patterns could carry functional costs.

Building on our initial findings in left-handed participants, we conducted the first-ever population-based study of hemispheric asymmetries using a stratified sample representative of the Flemish adult population. This study uniquely characterizes lateralization phenotypes across multiple functions and investigates their relationship to cognitive performance in a large sample (N = 200) drawn from the general population. We assessed hemispheric dominance for language, praxis, spatial attention, and face perception using fMRI tasks representative of each respective function. Participants also completed two behavioral test batteries assessing general cognitive ability and intelligence. We intentionally oversampled left-handed individuals to capture the full spectrum of hemispheric organization^32^. This design allowed us to (i) determine and map each participant’s pattern of hemispheric functional segregation, (ii) compare these patterns of functional segregation across handedness subgroups, and (iii) evaluate their behavioral relevance by linking them to neurocognitive performance.

## Results

A total of 200 individuals were included in the final sample, with equal representation of left-handed (*n* = 100) and right-handed (*n* = 100) participants and a balanced sex distribution (50 left-handed and 50 right-handed females) within groups. The sample composition was designed to closely represent the Flemish population in terms of sex, age, and educational level. See Figure 1 for a comparison of our sample with the population from which it was drawn. The left-handed participants ranged in age from 20 to 65 years (*M* = 41.08, *SD* = 13.14) and had a mean Edinburgh Handedness Inventory (EHI)^33^ score of −71.81 (*SD* = 32.81). The right-handed participants ranged in age from 19 to 65 years (*M* = 41.93, *SD* = 12.91) and had a mean EHI score of 93.6 (*SD* = 14.81). The handedness subgroups did not differ significantly in terms of age (*p* = .645), sex (*p* = 1), or years of education (*p* = .521).

**Figure 1.**
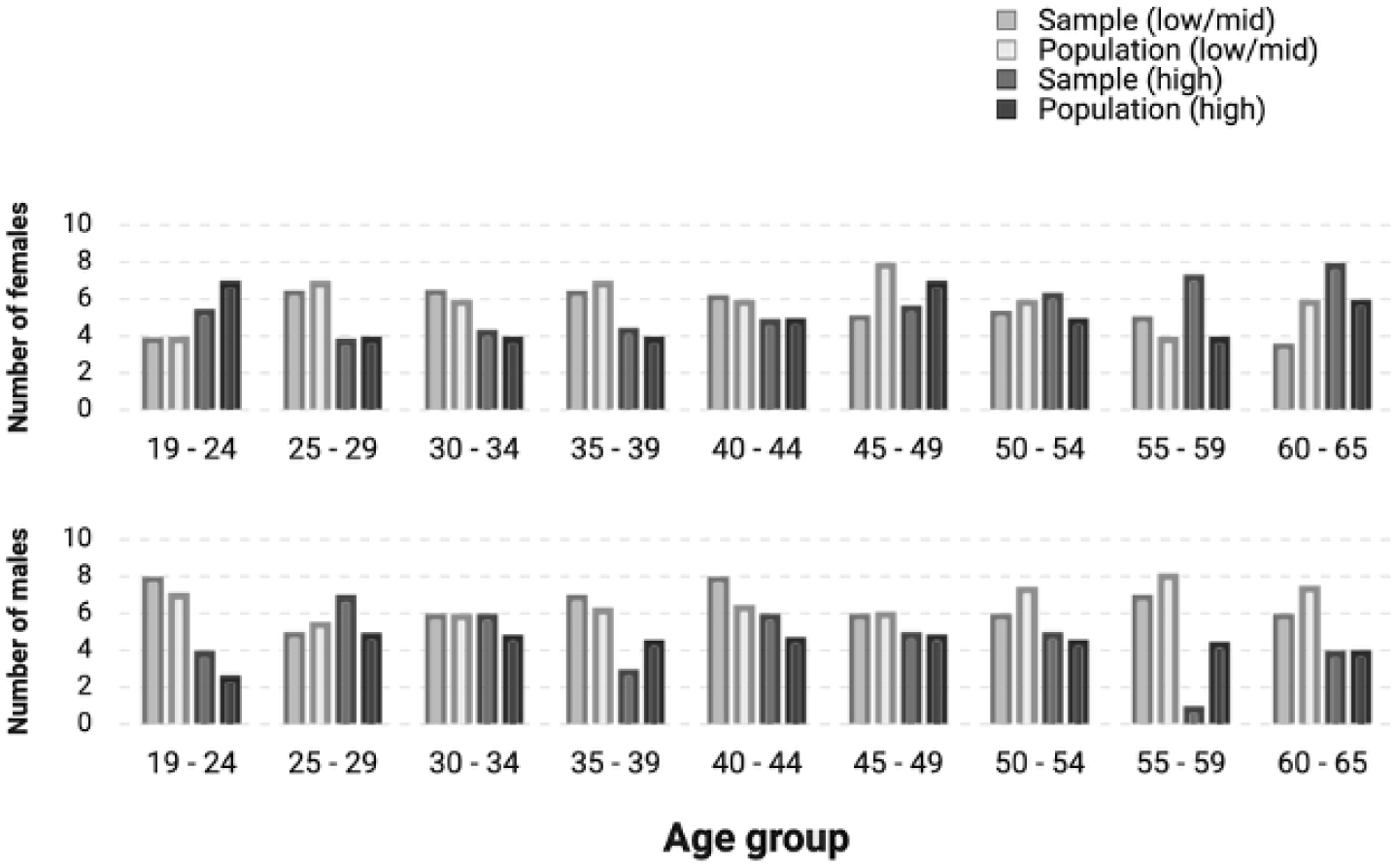
Comparison of our sample with the population from which it was drawn. Participants were recruited to match the age, sex, and educational-level distribution of the source population. Bars show the number of individuals in each category for the population and the sample, with education level in brackets and subdivided by sex. The sample closely mirrors population trends, with minor deviations in a few age groups.

Participant-specific laterality indices (LIs) were calculated for each of the four fMRI tasks to quantify hemispheric differences in activation. The LIs ranged from −1 to +1, with negative values indicating greater left hemisphere activity and positive values indicating greater right hemisphere activity^34^. The LIs were determined within regions of interest (ROIs) identified as crucial for the investigated functions, based on evidence from patient lesion studies (see Figure 2, panel 2; and Method for more detailed information).

**Figure 2.**
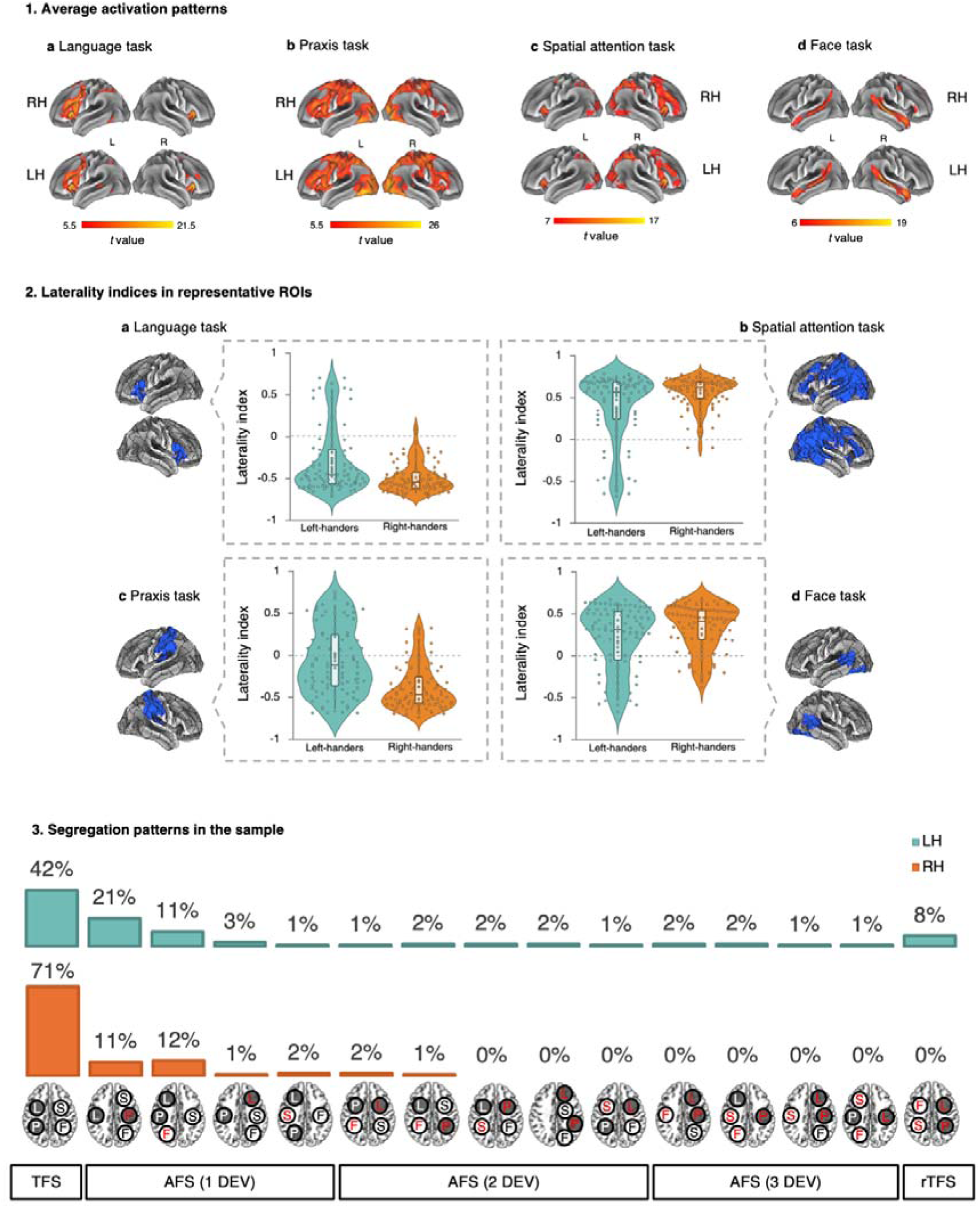
Summary of brain asymmetry data, including group-averaged values, individual laterality indices, and functional segregation patterns. Panel 1: average activation patterns for (**a**) language, (**b**) praxis, (**c**) spatial attention, and (**d**) face perception, by handedness; all FWE-corrected at pC<C.05, clusters >100 voxels. RH = right-handers, LH = left-handers, L = left hemisphere, R = right hemisphere. Panel 2: ROIs and violin plots showing individual LI values within each ROI for (**a**) language, (**b**) spatial attention, (**c**) praxis, and (**d**) face perception. Negative LI values represent relatively more left hemisphere processing, and positive LI values represent relatively more right hemisphere processing. Panel 3: shows an overview of hemispheric segregation phenotypes by handedness. Percentages indicate the proportion of individuals in each group with a given brain pattern. Red highlights indicate deviations from typical lateralization. LC=Clanguage, PC=Cpraxis, SC=Cspatial attention, FC=Cface perception; TFSC=Ctypical functional segregation, AFSC=Catypical functional segregation, rTFSC=Creversed TFS, DEVC=Cnumber of deviations from TFS.

### The relationship between functional asymmetries and handedness

We first examined the prevalence of right or left hemispheric dominance within the sample of left-handers (*n* = 100) and right-handers (*n* = 100). To calculate the percentage of individuals with a left or right hemispheric bias, participants were classified based on their individual LIs, using zero as the cut-off (Figure 2, panel 2; for a demonstration of different cut-off categorizations, see Supplementary Table 1). The percentages of hemispheric dominance for the four functions are presented in Table 1. Both handedness groups had significant population-level asymmetries: language and praxis were left-lateralized, while spatial attention and face perception were right-lateralized (see Table 1).

**Table 1.** Percentage of left- and right-handed individuals classified as left- or right-hemisphere dominant, with a comparison to 50% (no population-level bias).

| Function | Handedness group | Left hemisphere | Right hemisphere | $\chi^2$ | $p$-value | 95% CI's |
| --- | --- | --- | --- | --- | --- | --- |
| Language | Left-handers | **81%** | 19% | 37.21 | < 0.001 | 72%-88% |
|  | Right-handers | **97%** | 3% | 86.49 | < 0.001 | 92%-99% |
| Praxis | Left-handers | **60%** | 40% | 3.61 | 0.029 | 50%-69% |
|  | Right-handers | **88%** | 12% | 56.25 | < 0.001 | 80%-93% |
| Spatial attention | Left-handers | 16% | **84%** | 44.89 | < 0.001 | 76%-90% |
|  | Right-handers | 2% | **98%** | 90.25 | < 0.001 | 93%-99% |
| Face perception | Left-handers | 27% | **73%** | 20.25 | < 0.001 | 64%-81% |
|  | Right-handers | 15% | **85%** | 47.61 | < 0.001 | 77%-91% |

A series of χ² tests also revealed significant between-group differences in the distribution of hemispheric dominance for the two handedness groups. Compared with right-handers, left-handed participants were approximately 7.6 times more likely to be right hemisphere dominant for language [χ²(1) = 8.00; *p* = 0.005, 95% CI of the difference: 0.04, 0.23] and 4.89 times more likely to be right hemisphere dominant for praxis [χ²(1) = 18.95; *p* < 0.001, 95% CI of the difference: 0.15, 0.41]. Left-handed participants were also approximately 9.33 times more likely to be left hemisphere dominant for spatial attention compared to right-handed participants [χ²(1) = 10.32; *p* = 0.001, 95% CI of the difference: 0.05, 0.23]. However, there was no significant difference in the distribution of hemispheric dominance rates for face perception [χ²(1) = 3.65; *p* = 0.056]. Together, these findings indicate that left-handers show the same population bias as right-handers, although to a significantly lesser degree, except in face perception, where the bias is similar for both groups.

Second, we examined whether the handedness groups were significantly lateralized for the four different tasks using the calculated LIs (see Figure 2, panel 2). Left-handers were significantly lateralized for language (*m* = −0.30, *SD* = 0.38, t(99) = −7.75, *p* < 0.001, 95% CI [−0.37, −0.22]), spatial attention (*m* = 0.38, *SD* = 0.42, t(99) = 9.11, *p* < 0.001, 95% CI [0.30, 0.46]), and face perception (*m* = 0.21, *SD* = 0.38, t(99) = 5.49, *p* < 0.001, 95% CI [0.13, 0.28]), but not for praxis (*m* = 0.06, *SD* = 0.40, t(99) = −1.54, *p* = 0.506, 95% CI [−0.14, 0.02]). Right-handers were also significantly lateralized for language (*m* = −0.49, *SD* = 0.18, t(99) = −26.64, *p* < 0.001, 95% CI [−0.53, −0.45]), praxis (*m* = −0.39, *SD* = 0.27, t(99) = −14.39, *p* < 0.001, 95% CI [−0.44, −0.33]), spatial attention (*m* = 0.56, *SD* = 0.19, t(99) = 29.86, *p* < 0.001, 95% CI [0.52, 0.60]), and face perception (*m* = 0.32, *SD* = 0.28, t(99) = 11.54, *p* < 0.001, 95% CI [0.27, 0.38]).

### Hemispheric Functional Segregation in Individuals

To investigate individual patterns of hemispheric functional segregation, we combined each participant’s classification across the four tasks to create an overall categorization. The segregation patterns for individuals in both handedness groups are shown in Figure 2, panel 3. Since four distinct functional asymmetries were measured, the total number of possible combinations was 16 (2_). Remarkably, 15 out of these 16 combinations were observed (94%). The only pattern absent across all individuals was having all four functions lateralized to the left hemisphere.

Individuals were classified as having typical functional segregation (TFS) if language and praxis were lateralized to the left hemisphere, while spatial attention and face perception were lateralized to the right hemisphere. This pattern was the most common in both handedness groups, observed in 42% of left-handers and 71% of right-handers. Notably, 8% of left-handers and none of the right-handers had a complete reversal of TFS (referred to here as rTFS). Among the remaining participants with atypical functional segregation (AFS), 36% of left-handers and 26% of right-handers had one deviating function, while 14% of left-handers and 3% of right-handers had two or more deviating functions.

A χ² test indicated that the distribution of TFS, AFS, and rTFS differed significantly between the two handedness groups [χ²(2, *N* = 200) = 21.03, *p* < 0.001]. Post hoc tests with Bonferroni correction revealed significant differences for TFS with higher rates in right-handers (71%) compared to left-handers (42%) (*p* < 0.001), AFS was higher in left-handers (50%) compared to right-handers (29%) (*p* = 0.014), and rTFS was higher in left-handers (8%) compared to right-handers (0%) (*p* = 0.023).

### Hemispheric Functional Segregation and Neurocognitive Performance

Next, we investigated whether deviation from prototypical hemispheric segregation influenced neurocognitive performance. To assess this, we used a general cognitive test battery (Repeatable Battery for the Assessment of Neuropsychological Status [RBANS]) and measures of intelligence using standardized intelligence quotients (IQ; Wechsler Abbreviated Scale of Intelligence [WASI II] or Raven’s progressive matrices [RPM]). Both tests are reported as standard scores with a mean of 100 and a standard deviation of 15. To ensure that handedness did not confound the results, we first compared right- and left-handers. There was no significant difference in total RBANS scores between the two groups (*M* left-handers = 98.59, *SD* = 12.62; *M* right-handers = 99.79, *SD* = 12.42), *t*(197.95) = 0.68, *p* = .499, d = 0.10. Similarly, no significant difference was found in IQ scores (*M* left-handers = 102.47, *SD* = 15.45; *M* right-handers = 105.48, *SD* = 15.50), *t*(198) = −1.38, *p* = .171, d = 0.20. Note that the average performance scores and standard deviations of all subgroups are very close to the preset standards, indicating that the sample is representative of its population (see Figure 3).

**Figure 3.**
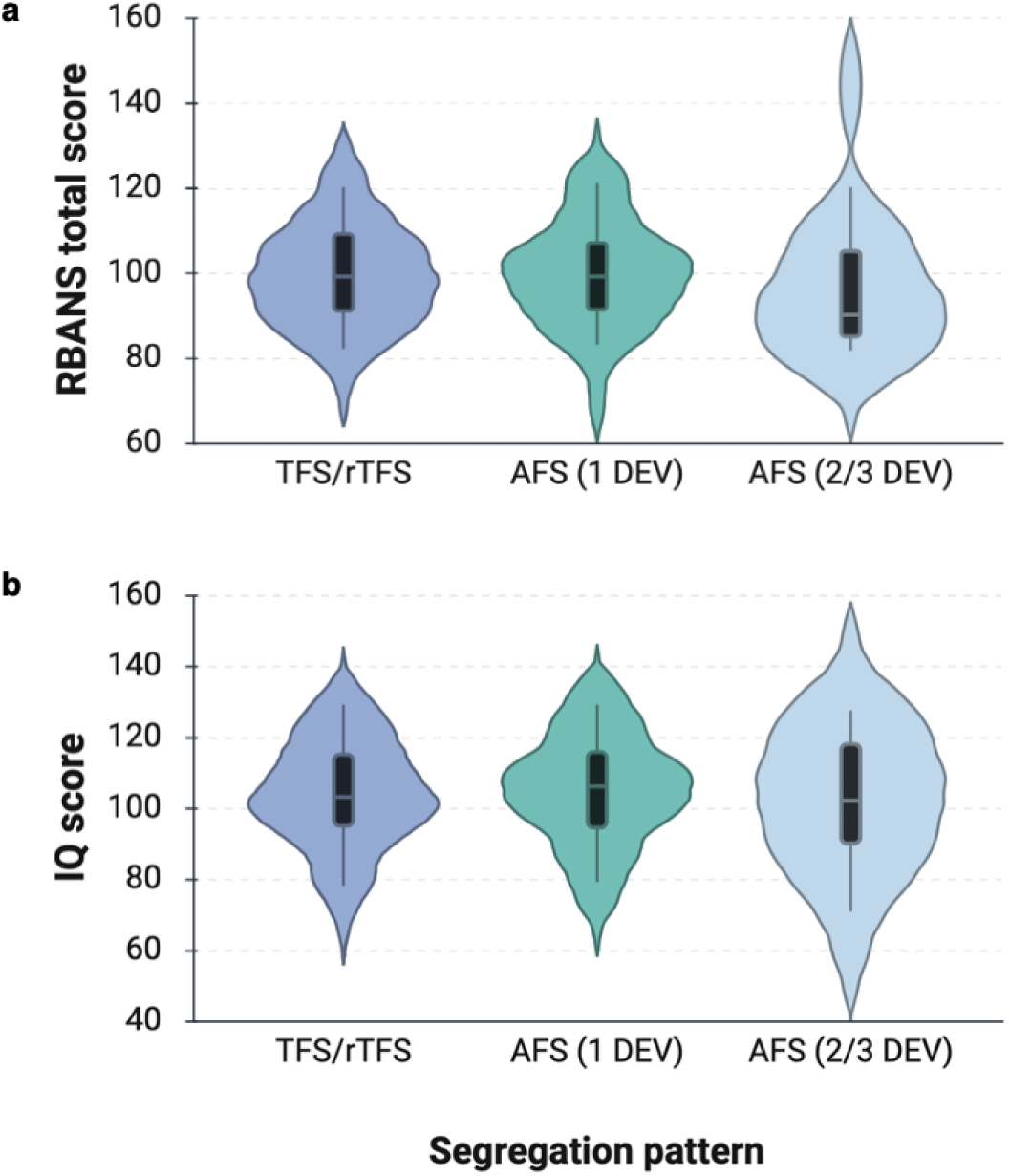
Neurocognitive performance across functional segregation subgroups. Violin plots show the distribution of **a** RBANS total scores and **b** IQ scores. Violin plots show the data distribution; boxes indicate median and IQR, whiskers the 5th–95th percentiles Participants were classified based on the number of deviations from typical functional segregation: typical/reversed typical functional segregation (TFS/rTFS, n = 121), one deviation (AFS 1 DEV, n = 62), and two or three deviations (AFS 2/3 DEV, n = 17).

We then examined whether the identified patterns of functional segregation were associated with neurocognitive performance. To this end, participants were divided into three subgroups: (1) typical functional segregation (*n* = 113) or reversed typical functional segregation (*n* = 8), (2) one deviation (*n* = 62), and (3) two or three deviations (*n* = 17) from typical functional segregation. The grouping of TFS and rTFS was based on the assumption that typical hemispheric segregation is preserved across both phenotypes (just reversed in a subset). There were no significant differences in RBANS scores between the groups, *F*(2, 197) = 0.35, *p* = .704, ω^2^ = 0, nor in IQ scores, *F*(2, 197) = 0.24, *p* = .788, ω^2^ = 0.

In a second step, we investigated whether the extent of an individual’s deviation from the ‘prototypical’ TFS pattern was associated with cognitive performance. We first determined the mean LI for each task in the right-handed TFS group (i.e., “textbook” brains; *n* = 71) and used these averages as a reference. For each participant, we then calculated the extent to which their LI differed from this reference for each task (taking the LI’s sign into account) and summed the absolute differences to obtain a TFS-Deviation score. For transparency, we excluded the eight left-handed participants with rTFS. Pearson correlations revealed no significant association between the TFS-Deviation score and total RBANS score (*r* = –.052, *p* = .471) or IQ score (*r* = –.026, *p* = .722). As a last step, we also compared the 20 highest-performing individuals (total RBANS scores > 118) with the 20 lowest-performing individuals (total RBANS scores < 84) in terms of handedness, LIs for the four tasks, number of atypically lateralized functions, segregation pattern (TFS, AFS, rTFS), and TFS-Deviation score. No significant group differences were observed.

## Discussion

Our findings confirm and replicate established population-level biases: language and praxis are typically left-lateralized^2,4,35^, while spatial attention and face processing tend to engage the right hemisphere^6–9^. They also extend the well-established higher prevalence of atypical hemisphere dominance in left-handers, traditionally based on single-function studies, by demonstrating that deviations from the typical pattern of hemispheric segregation are the norm rather than the exception in left-handed individuals. Notably, our results demonstrate that such deviations are not restricted to left-handers; over one quarter of right-handed participants had at least one atypically lateralized function. Crucially, these deviations were not associated with diminished cognitive performance or lower intelligence, suggesting that, at least in a healthy sample, such variability occurs naturally without necessitating an underlying pathology—or even a minor cognitive disadvantage—as hypothesized by earlier theories of hemispheric dominance.

### Reappraising the prototypicality of hemispheric segregation

Our study is the first to demonstrate the full phenotypic range of hemispheric functional segregation across four asymmetrical cognitive functions in a stratified sample drawn from the general population, albeit with a deliberate overrepresentation of left-handed individuals. The most common pattern of hemispheric functional segregation was the prototypical one, with language and praxis lateralized to the left hemisphere and spatial attention and face processing lateralized to the right hemisphere. This pattern was observed in about 70% of right-handers and 40% of left-handers. While these findings suggest a general bias toward maintaining typical hemispheric functional segregation, this bias is not absolute and allows for individual variation. This variation was particularly pronounced in left-handed individuals, 58% of whom deviated from the prototypical directional pattern. This suggests that when one function shows atypical lateralization (such as left-handedness, if handedness were also used as a function), there is a higher likelihood that other functions will also be atypically lateralized. Left-handedness, therefore, emerges as a predictor of atypical functional segregation (AFS). However, notable variation was also observed in right-handers, with 29% showing deviations from the typical pattern.

Another key finding was that a complete reversal of hemispheric functional segregation occurred exclusively in left-handed individuals. There are currently two main theoretical accounts of the relationship between lateralized functions. Causal accounts propose that the lateralization of one function to one hemisphere causes, in some fashion, another function to lateralize to the opposite hemisphere^21,36–39^. Although accounts differ in the specific mechanisms underlying this event, they propose that cognitive functions do not lateralize independently of one another. In contrast, statistical accounts^21,22^ assume that whatever underlies the lateralization of a function reflects independent probabilistic biases. Our findings challenge causal accounts, which cannot explain both functional crowding and the dissociations observed among the four measured functions. Simultaneously, statistical accounts are insufficient to explain the relatively high prevalence of left-handers with reversed patterns of functional segregation (8%) if functions lateralize independently. Collectively, these results challenge existing interpretations and suggest the need for a refined model that extends the statistical hypothesis to accommodate the observed patterns of hemispheric organization.

Previously, we have proposed the existence of a segregation bias^30,40^, which guides the brain’s development according to a predefined blueprint for organizing functions that may be genetically determined. For most individuals, this blueprint codes for the typical pattern of functional segregation (TFS). This process likely emerges early in embryonic development, akin to the one that establishes left–right asymmetry in visceral organs^41^. Crucially, this segregation bias does not determine lateralization of each function independently; rather, it reflects a higher-order plan for organizing multiple functions across the hemispheres. Occasionally, this blueprint is reversed, giving rise to individuals with a mirror-reversed configuration but an equally structured segregation pattern (rTFS). Once this global template is in place, the lateralization of individual functions may still be shaped by probabilistic mechanisms, consistent with the statistical hypothesis^22,42^. As a result, even when the segregation bias is predetermined, whether typical or reversed, deviations from prototypical or mirrored patterns will occur^40^, as observed in our study. This framework also helps explain why left-handers show greater variability in hemispheric asymmetries: If handedness is a modular trait that is also reversed in individuals with a reversed blueprint, some of this increased variability may reflect the existence of a reversed blueprint, while other instances of atypical lateralization may arise from weaker genetic influences or from ontogenetic factors acting on function-specific development.

### Hemispheric functional segregation is not associated with general cognitive ability

In contrast to what some evolutionary models would predict based on the premise that prototypical functional segregation is the most optimal division of labor nature could generate^23,25,26^, our results do not indicate that atypical hemispheric division leads to marked cognitive disadvantages in terms of general cognitive ability or intelligence. While prototypical hemispheric organization may reflect common population-level patterns, it does not appear to confer a clear cognitive advantage. No significant differences were observed between participants with a prototypical configuration, those with one functional/modular deviation, and those with multiple deviations. Although this finding contradicts a smaller study conducted in our lab, which examined functional segregation in left-handers with right- and left-hemisphere dominance for language^30^, it reassuringly suggests that general cognitive ability is not necessarily better or worse depending on how hemispheric asymmetries are organized. Importantly, our results should not be taken to imply that variations in segregation patterns, or better, variations in individual asymmetries, completely lack consequences for behavior. Our cognitive battery may not have been sensitive enough to capture more subtle differences. It could well be that the effects of segregation patterns are more evident in domain-specific cognitive processes^11,32^. For example, McManus^11^ proposes that talents and deficits arising from atypical inter- and intra-hemispheric modular connections are closely linked to the specific asymmetries involved. One could, for instance, imagine that someone exceptionally skilled in a given domain might possess a corresponding hemispheric configuration that supports that talent. However, there is currently no empirical evidence supporting this link between talent and asymmetry.

On the other end of the spectrum, findings suggest that certain configurations of functional lateralization may be disadvantageous under specific cognitive demands. For example, Zhu and Cai^43^ found evidence that individuals with language and spatial attention lateralized to the same hemisphere performed worse on dual tasks involving both functions than those with a more distributed hemispheric organization. A review of asymmetry in language and spatial attention lateralization found that individuals with both functions in the same hemisphere performed worse on tasks such as language comprehension, spatial skills, and reading^12^. This suggests that overlapping lateralized functions may interfere with one another when co-activation is required, potentially leading to performance costs.

### Distinguishing adaptive from maladaptive patterns of asymmetry

Our study highlights the flexibility with which functions can be distributed across the hemispheres, with seemingly little to no cost to behavior. This apparent flexibility is somewhat at odds with the higher prevalence of unusual asymmetry so often reported in neurodevelopmental and psychiatric conditions^14^, and highlights the need to better understand the mechanisms that govern hemispheric segregation. Of course, it is essential to note that most of these clinical studies have focused on the *group average*, and the individual variability in asymmetry patterns for many of them remains to be established. For example, we recently provided evidence suggesting that developmental dyslexia may be linked to inconsistent lateralization within sub-regions of the phonological and lexical system^44^, rather than a uniform reduction in asymmetry across the entire sample. A similar trend was observed across language tasks in individuals with various language impairments^45^. Thus, several outstanding questions remain as to when variability in asymmetry may turn maladaptive. We argue that addressing these questions requires careful attention to individual differences, as understanding the full spectrum of asymmetry patterns is essential for our fundamental understanding of brain organization and for relevant clinical applications.

### Beyond group averages: the value of individual differences

More broadly, our findings underscore that individual differences are not merely statistical noise but instead provide meaningful and essential insights into the organizing principles of the human brain. Examining variability at the individual level reveals intricate patterns obscured by group averages and challenges conventional assumptions about hemispheric specialization. Recognizing this variability is essential not only for refining theoretical models of hemispheric specialization but also for advancing personalized diagnosis and rehabilitation in neuropsychology and neurology. Importantly, these differences should inform the development of lateralized treatment approaches, such as neurostimulation protocols, which often rely on standard assumptions of functional asymmetry. Embracing individual variability represents a crucial step toward more precise, personalized clinical care and a deeper understanding of brain function and its behavioral consequences.

## Method

### Participants

A total of 202 individuals were recruited via a research website, social media, word of mouth, flyers displayed in local public spaces and businesses, and advertisements during local public science events. Inclusion criteria were native-level proficiency in Dutch and no history of brain injury or neurological disease. Recruitment efforts specifically targeted left-handed individuals to ensure a sufficiently large sample of this group. Participants received 50 euros as reimbursement and compensation for parking or public transport costs.

Sample stratification was based on 2021 demographic data from StatBel (Algemene Directie Statistiek – Statistics Belgium), reflecting the active and inactive populations in Flanders. The sample was stratified by biological sex, age (nine age groups spanning 19-65), and level of education (short to mid: International Standard Classification of Education (ISCED) 1-4; long: ISCED 5-8). Demographic counts were proportioned into 36 cells (2 sex × 2 education × 9 age intervals) with corresponding target sample sizes (see Figure 1). With few exceptions, cells were adequately covered, supporting the representativeness of the sample for the typical Flemish (Western European) population.

Two participants (one left-handed and one right-handed, both male) were excluded from the analysis due to excessive head movement during fMRI scanning (> 7 mm movement within at least one of the functional scans), resulting in a final sample of 200 (100 left-handed, 100 right-handed), balanced for sex (50 females per handedness group). Left-handed participants were aged 20 to 65 years (*M* = 41.08, *SD* = 13.14) with a mean Edinburgh Handedness Inventory (EHI) score of −71.81 (*SD* = 32.81). The right-handed participants were aged 19 to 65 years (*M* = 41.93, *SD* = 12.91) and had a mean EHI score of 93.6 (*SD* = 14.81). The handedness subgroups did not differ significantly in age (*P* = .645), sex (*P* = 1), or years of education (*P* = .521). The study was approved by the Medical Ethics Committee of Ghent University Hospital (approval number ONZ-2022−0118), and written informed consent was obtained from all participants.

## Stimuli, Materials, and Procedures

### Behavioral measures

All participants completed a handedness questionnaire, a behavioral battery of tests to assess general cognitive ability, and an intelligence test.

Hand preference was assessed using a Dutch version of the Edinburgh Handedness Inventory (EHI)^33^. This questionnaire asks individuals to report their hand preference for ten actions involving hand use, such as writing, throwing, and striking a match. Participants indicate whether they ‘always’ or ‘usually’ perform each action with their left or right hand, or whether they have no preference. Based on these responses, a hand preference score is calculated, ranging from −100 (indicating complete left-hand preference) to +100 (indicating complete right-hand preference).

General cognitive performance was assessed using a Dutch version of the Repeatable Battery for the Assessment of Neuropsychological Status (RBANS)^46^. The RBANS takes approximately 30 minutes to complete and consists of 10 cognitive subtests that contribute to five index scores: Immediate Memory (based on a 10-word List Learning and Story Memory subtest), Visuospatial/Constructional (based on a Complex Figure copy and Line Orientation subtest), Language (based on a Picture Naming and Semantic Fluency subtest), Attention (based on a Digit Span and Symbol Digit Coding subtest), and Delayed Memory (based on delayed recall of List Learning, Story Memory, and Complex Figure performance). Performance on these subtests was combined into a Total Scale score, reported as a standardized score with a mean of 100 and a standard deviation of 15.

Intelligence was assessed using the Dutch version of the Wechsler Abbreviated Scale of Intelligence, Second Edition (WASI-II)^47^, or Raven’s Progressive Matrices (RPM^48^. A total of 162 individuals had their IQ assessed with the WASI-II, and 38 with the RPM. The two-subtest version of the Wechsler Abbreviated Scale of Intelligence, consisting of Vocabulary and Matrix Reasoning, was used and can be administered in approximately 15 minutes. The Vocabulary subtest measures an individual’s breadth of verbal knowledge, concept formation, and expressive language skills. It includes 31 items that require participants to define the presented concepts. The Matrix Reasoning subtest assesses nonverbal abstract reasoning, problem-solving, and perceptual organization skills. It consists of 30 visual pattern series, each with one missing element, and the participant must select the correct piece from a set of options to complete the pattern. The RPM is a multiple-choice test of nonverbal abstract reasoning. It consists of 60 progressively difficult items, each a 3 × 3 matrix of geometric designs with one missing element. Participants select the design that completes the pattern from a set of six to eight choices.

### fMRI paradigm

Participants completed four fMRI tasks to obtain participant-specific measures of lateralization for language (word generation task), praxis (tool pantomiming task), spatial attention (line bisection judgment task), and face perception (dynamic faces one-back task). Example stimuli are shown in Supplementary Figure 1. All tasks were programmed and run using PsychoPy, version 2022.2.4.

### Verbal fluency task

Language lateralization was assessed using a covert single-letter phonemic verbal fluency task in a blocked design^49^. The task consisted of seven experimental blocks, seven control blocks, and 14 rest blocks, each lasting 15 seconds. During the experimental blocks, participants covertly generated as many words as possible, starting with a letter presented in the center of the screen. Seven letters (b, d, k, m, p, r, and s) were used, one per experimental block. In the control blocks, participants covertly repeated the meaningless string “baba” displayed in the center of the screen. During the rest blocks, a fixation cross appeared, and participants were instructed to relax and not think of anything.

### Tool pantomime task

Praxis lateralization was assessed using a tool pantomiming task in a blocked design^5,35^. The task consisted of 12 experimental blocks and 12 control blocks, each lasting 21 seconds and separated by 15-second rest blocks. In both the experimental and control blocks, the stimuli consisted of two object pictures, one presented on the left and the other on the right side of the screen. In the experimental blocks, participants pantomimed using familiar tool pairs with their hands, such as a pencil and a sharpener, according to their positions on the screen. For example, if a pencil appeared on the left and a sharpener on the right, participants pretended to use their left hand to hold the pencil while sharpening it with their right hand. There were 20 unique tool pairs, half with the active object on the right and half on the left, resulting in 40 tool combinations. In the control blocks, participants performed a bimanual rotation movement in response to simple object cues, with 12 unique rotation stimuli (6 with the right hand and 6 with the left). All experimental and control blocks consisted of six items, each presented for 3.5 seconds. Participants were instructed to perform pantomimes calmly, moving only their hands and wrists while keeping their arms and heads still to minimize motion artifacts. During the rest blocks, a fixation cross was shown in the center of the screen, and participants were instructed to relax and not think of anything.

### Dynamic faces task

Face lateralization was measured using a dynamic faces task in a blocked design^50^. The task consisted of eight experimental blocks and eight control blocks, separated by 12-second rest blocks. In the experimental blocks, participants watched video clips of faces that displayed dynamic changes in expression, transitioning from neutral to happy or from neutral to sad. A total of 40 unique video clips were presented in a pseudorandom order. In the control blocks, participants viewed clips of inanimate objects undergoing subtle movements without significant positional changes. A total of 40 unique video clips were also presented in a pseudorandom order. Each block contained six 2-second clips, with one clip being an identical repetition of a previous clip. Participants were required to press buttons with both hands whenever a repetition occurred. During the rest blocks, a fixation cross was shown in the center of the screen, and participants were instructed to relax and not think of anything.

### Landmark task

Spatial attention lateralization was assessed using a landmark task in a blocked design^51^. The task consisted of six experimental blocks and six control blocks, each lasting 21.6 seconds and separated by 15-second rest blocks. In the experimental blocks, participants viewed a horizontal black line with a short vertical mark that was either centered or offset by 2.5%, 5.0%, or 7.5% of the line’s length to the left or right. Participants were instructed to press response buttons with their left and right index fingers simultaneously when the mark appeared precisely at the center of the line. The control blocks used the same horizontal stimuli, but the vertical mark was either slightly above or in contact with the horizontal line. Participants pressed both buttons only if the mark touched the line. A four-second instruction screen preceded all experimental and control blocks. During each block, a total of 12 different stimuli were presented for 1.8 seconds each. There are seven experimental stimuli in total: three bisected to the left, three bisected to the right, and one bisected in the middle. In addition, there were 14 control stimuli, comprising identical versions of the experimental stimuli and corresponding versions in which the vertical mark did not touch the horizontal line. During the rest blocks, a fixation cross was shown in the center of the screen, and participants were instructed to relax and not think of anything.

### MRI processing

#### MRI acquisition

The scans were acquired using a Siemens 3 Tesla Prisma magnetic resonance (MR) scanner located at Ghent University Hospital, equipped with a 64-channel head coil. Functional images were obtained with a T2-weighted gradient-echo EPI sequence (multiband factor: 4) with the following parameters: repetition time (TR) = 1070 ms, echo time (TE) = 31 ms, flip angle (FA) = 52°, field of view (FOV) = 210 x 210 mm, 60 slices; acquired voxel size = 2.5 x 2.5 x 2.5 mm (reconstructed voxel size = 2.5 x 2.5 x 2.5 mm). The number of acquisition volumes for each task was as follows: 393 for the word-generation task, 556 for the tool-pantomiming task, 362 for the dynamic-faces task, and 456 for the landmark task. An additional five initial volumes for each functional run were acquired and subsequently discarded to establish steady-state magnetization.

T1-weighted structural images were obtained using an MPRAGE sequence with the following scan parameters: TR = 2250 ms, TE = 4.18 ms, inversion time (TI) = 900 ms, FA = 9°, FOV = 256 x 256 mm, 176 slices, voxel size = 1 x 1 x 1 mm (reconstructed voxel size = 1 mm³), and an acquisition time of 314 seconds.

### MRI preprocessing

Preprocessing was performed using *fMRIPrep* 21.0.2^52^ which is based on *Nipype* 1.6.1^53^.

#### Anatomical data preprocessing

The T1-weighted (T1w) image was corrected for intensity non-uniformity (INU) with N4BiasFieldCorrection, distributed with ANTs 2.3.3^54^, and used as T1w-reference. The T1w-reference was skull-stripped using a *Nipype* implementation of the antsBrainExtraction.sh workflow (from ANTs), with OASIS30ANTs as the target template. Brain tissue segmentation of cerebrospinal fluid (CSF), white-matter (WM), and grey-matter (GM) was performed on the brain-extracted T1w using fast ^55^ (FSL). Volume-based spatial normalization to standard space (MNI) was performed through nonlinear registration with antsRegistration (ANTs 2.3.3), using brain-extracted versions of both the T1w reference and the T1w template. FSLs *MNI ICBM 152 non-linear 6th Generation Asymmetric Average Brain Stereotaxic Registration Model*^56^ was used for spatial normalization.

#### Functional data preprocessing

For each of the four BOLD runs, the following preprocessing steps were performed. First, a reference volume was generated using a custom fMRIPrep methodology for head motion correction. Head motion parameters, including transformation matrices and six corresponding rotation and translation parameters, were estimated prior to spatiotemporal filtering using mcflirt (FSL)^57^. The BOLD time series were resampled into their original native space by applying transformations to correct for head motion. The BOLD reference was co-registered to the T1w reference using mri_coreg (FreeSurfer), followed by flirt (FSL 6.0.5.1:57b01774) ^58^with the boundary-based registration cost function^59^. Co-registration was configured with six degrees of freedom. Both a reference volume and its skull-stripped version were generated using fMRIPrep’s custom methodology. Automatic removal of motion artifacts was performed using independent component analysis (ICA-AROMA)^60^ on the preprocessed BOLD time series in MNI space, after removal of non-steady-state volumes and spatial smoothing with an isotropic Gaussian kernel of 6 mm full-width at half-maximum (FWHM). Corresponding “non-aggressively” denoised runs were produced following this smoothing. Gridded (volumetric) resamplings were performed using antsApplyTransforms (ANTs), configured with Lanczos interpolation to minimize smoothing effects^61^. Non-gridded (surface) resamplings were performed using mri_vol2surf (FreeSurfer).

#### fMRI analysis

After preprocessing in fMRIPrep, analysis was continued in SPM12 running in MATLAB_R2021B. In SPM12, the general linear model was used to predict the hemodynamic response curve by applying regressors that coded for each experimental and control condition, along with the six movement parameters (three rotation and three translation) estimated during preprocessing. A boxcar function, convolved with the hemodynamic response function, was fitted to the time series at each voxel, resulting in weighted beta images. The beta images of the experimental and control conditions were then contrasted to create a t-statistic map.

#### Determination of Participant-Specific lateralization

The LI-toolbox plugin for SPM^62,63^ was used to assess hemispheric contribution based on the t-statistic map for each task. This toolbox allows comparisons between the right and left hemispheres while addressing common issues such as statistical outliers, threshold-dependent comparisons, and data sparsity. It employs a bootstrapping method to calculate a laterality index (LI) ranging from −1 to +1 using the standard formula: LI = (L - R) / (L + R). To align with the Laterality Indices Consensus Initiative (LICI)^34^, the scores were inverted, so that negative values indicate greater left hemisphere activation and positive values indicate greater right hemisphere activation. Participant-specific LIs were calculated for task-specific regions of interest (ROIs; see below) for each of the four fMRI tasks, using the recommended settings and the weighted mean LI calculation method.

#### Regions of interest

ROIs were calculated for each fMRI task using the HCP-MMP1 atlas^64^ focusing on areas whose damage, as indicated by lesion studies, disrupts the investigated function^65–72^. Each ROI was made symmetrical by selecting the left and right ROI, flipping them over the x-axis, and then merging them. The ROIs are visualized in Figure 2.

#### Statistical analyses

All statistical analyses were conducted using R version 4.2.2 in RStudio 2023.06.1+524. An alpha level of p < 0.05 was used to determine statistical significance for all tests. Bonferroni corrections were applied where multiple comparisons were performed.

#### Demographics

To assess differences in demographic variables (age, years of education, and handedness scores) between left- and right-handed participants, we used independent samples t-tests. Sex distribution was compared using a chi-square (χ²) test.

#### Laterality Indices and Functional Segregation

Participant-specific Laterality Indices (LIs) were computed for each of the four fMRI tasks (language, praxis, spatial attention, and face perception) using the weighted mean method in the LI-toolbox. LIs ranged from −1 (strong left hemisphere dominance) to +1 (strong right hemisphere dominance). To classify participants as left- or right-hemisphere dominant for each function, we set the LI threshold to 0, categorizing values < 0 as left-dominant and values > 0 as right-dominant. The prevalence of left- and right-hemisphere dominance was compared against 50% using one-tailed Pearson’s chi-square (χ²) tests, as we predicted that the proportion would be larger than 50%. The distribution of hemispheric dominance was also compared between handedness groups using Pearson’s chi-square (χ²) tests. To determine whether the two handedness groups had significant population-level lateralization biases, one-sample t-tests were conducted against a population mean of 0 (indicating no hemispheric preference).

#### Patterns of Functional Segregation

To examine how functional asymmetry varied across individuals, we categorized participants into functional segregation groups based on their individual laterality profiles across all four fMRI tasks (language, praxis, spatial attention, and face perception). First, we identified the range of segregation patterns present in the dataset. To examine how functional asymmetry varied across individuals, we categorized each participant into one of three segregation patterns: Typical Functional Segregation (TFS), Atypical Functional Segregation (AFS), or Reversed Typical Functional Segregation (rTFS). The proportions of individuals in each handedness group were compared using a chi-square (χ²) test.

#### Functional Segregation and Neurocognitive Performance

To examine the relationship between hemispheric functional segregation and neurocognitive performance, we conducted a series of statistical comparisons. First, we assessed whether handedness itself influenced cognitive performance. Independent samples t-tests were performed to compare RBANS total scores and IQ scores between left- and right-handed participants. This analysis ensured that any observed effects in later comparisons were not confounded by differences in overall cognitive ability between the two handedness groups.

Next, we investigated whether deviations from the prototypical functional segregation pattern were associated with neurocognitive performance. Participants were classified into one of three functional segregation groups: (1) Typical Functional Segregation (TFS) – Language and praxis were left-lateralized, while spatial attention and face perception were right-lateralized, and Reversed Typical Functional Segregation (rTFS) – A mirror image of TFS, with language and praxis right-lateralized and spatial attention and face perception left-lateralized. Because few participants exhibited rTFS, this group was merged with the TFS group for analysis, assuming they display a prototypical, but reversed, pattern of functional segregation. (2) Atypical Functional Segregation (AFS-One Deviation) – Participants exhibited a single deviation from the typical pattern (e.g., right-lateralized language with typical lateralization for the other three functions). (3) Atypical Functional Segregation (AFS-Multiple Deviations) – Participants exhibited two or more deviations from the typical pattern, indicating a more widespread atypical lateralization profile.

To assess whether functional segregation patterns influenced cognitive performance, a one-way ANOVA was conducted, comparing RBANS total scores and IQ scores across the three groups: TFS + rTFS, AFS-one dev, and AFS-multiple dev, regardless of handedness. Post hoc Tukey’s tests were performed where significant differences were found.

We also investigated whether the extent of an individual’s deviation from the ‘prototypical’ TFS pattern was associated with cognitive performance. We first determined the mean LI for each task in the right-handed TFS group (i.e., “textbook” brains; *n* = 71) and used these averages as a reference. For each participant, we then calculated the extent to which their LI differed from this reference for each task (taking the LI’s sign into account) and summed the absolute differences to obtain a TFS-Deviation score. For transparency, we excluded the eight left-handed participants with rTFS. We then performed Pearson correlations between the resulting score and RBANS total scores and IQ scores, respectively.

We then compared the 20 highest-performing individuals (total RBANS scores > 118) with the 20 lowest-performing individuals (total RBANS scores < 84) in terms of handedness, LIs for the four tasks, number of atypically lateralized functions, segregation pattern (TFS, AFS, rTFS), and TFS-Deviation score, using independent samples t-tests.

#### Outlier Handling

Data points with LI scores beyond 2.5 standard deviations from the mean were identified as potential outliers. Based on this criterion, one participant had an outlier score on the RBANS battery. However, since this score reflects performance on a standardized test, we opted to include this participant. For transparency, the inclusion or exclusion of this participant does not affect the analysis’s outcome. The processed data and analysis scripts are available on OSF at https://osf.io/jvrzc/.

## Acknowledgements

Most graphs used in the figures were created in BioRender. Karlsson, E. (2025), including the bar graphs in Figure 1 (https://BioRender.com/g6ezmc3), the four violin plots in Figure 2 (https://BioRender.com/8z64q4n, https://BioRender.com/ktbfoy9, https://BioRender.com/vj2ixfz, https://BioRender.com/eqknp5v), and the RBANS graph (https://BioRender.com /9renwf4) and IQ graph (https://BioRender.com/g6g3opd) in Figure 3.

## Notes

This work was supported by a Ghent University Special Research Fund (BOF; BOF/24J/2021/172) grant assigned to Guy Vingerhoets.

https://osf.io/jvrzc/

### Competing Interest Statement

The authors have declared no competing interest.

