## Supplemental materials for "The individual brain: mapping variability in hemispheric functional organization across cognitive domains in a representative sample"

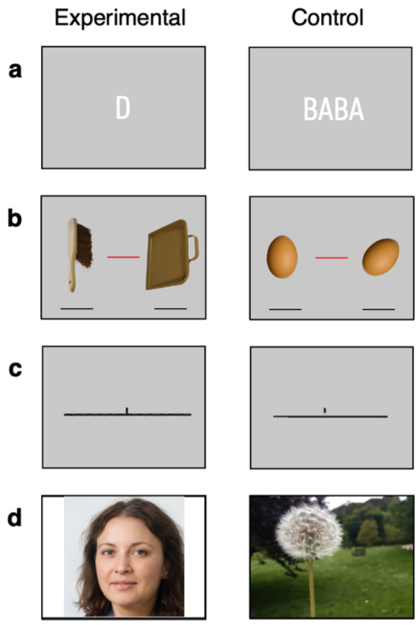


**Supplementary Figure 1.** *Example stimuli used in the block-design fMRI localizers. Panel* ***a*** *shows the experimental and control stimuli used in the verbal fluency (language) task,* ***b*** *in the praxis (tool-use) task,* ***c*** *in the landmark (spatial attention) task, and* ***d*** *in the face perception localizer (face shown is a computer-generated image via ThisPersonDoesNotExist of a person that does not exist but is representative of the stimuli used).*

| Task | Handedness group | LI cut-off ± 0 | | 0.1 | | | 0.2 | | | 0.3 | | | 0.4 | | |
|---|---|---|---|---|---|---|---|---|---|---|---|---|---|---|---|
| | | L | R | L | R | B | L | R | B | L | R | B | L | R | B |
| **Language** | **Left-handers** | 81 | 19 | 78 | 17 | 5 | 72 | 14 | 14 | 68 | 12 | 20 | 60 | 12 | 28 |
| | **Right-handers** | 97 | 3 | 95 | 1 | 4 | 91 | 0 | 9 | 85 | 0 | 15 | 78 | 0 | 22 |
| **Praxis** | **Left-handers** | 60 | 40 | 51 | 32 | 17 | 45 | 28 | 27 | 36 | 22 | 42 | 23 | 21 | 56 |
| | **Right-handers** | 88 | 12 | 85 | 8 | 7 | 76 | 5 | 19 | 73 | 3 | 24 | 62 | 0 | 38 |
| **Spatial attention** | **Left-handers** | 16 | 84 | 16 | 82 | 2 | 14 | 76 | 10 | 12 | 70 | 18 | 12 | 63 | 25 |
| | **Right-handers** | 2 | 98 | 2 | 96 | 2 | 0 | 94 | 6 | 0 | 90 | 10 | 0 | 84 | 16 |
| **Face perception** | **Left-handers** | 27 | 73 | 22 | 69 | 9 | 19 | 69 | 18 | 16 | 52 | 32 | 12 | 39 | 49 |
| | **Right-handers** | 15 | 85 | 13 | 78 | 9 | 8 | 72 | 20 | 2 | 63 | 35 | 0 | 51 | 49 |

Notes: L = left hemisphere category, R = right hemisphere category, B = bilateral category

**Supplementary Table 1.** *Hemispheric dominance classifications across different laterality index (LI) cut-offs for the two handedness groups, shown separately for each of the four cognitive tasks. Each number represents an individual. L = left hemisphere category, R = right hemisphere category, B = bilateral category.*
